# Home-field advantage affects the local adaptive interaction between *Andropogon gerardii* ecotypes and rhizobiome

**DOI:** 10.1101/2023.01.04.522809

**Authors:** Anna Kazarina, Soumyadev Sarkar, Shiva Thapa, Leah Heeren, Abigail Kamke, Kaitlyn Ward, Eli Hartung, Qinghong Ran, Matthew Galliart, Ari Jumpponen, Loretta Johnson, Sonny T.M. Lee

## Abstract

Due to climate change, drought frequencies and severities are predicted to increase across the United States. Plant responses and adaptation to stresses depend on plant genetic and environmental factors. Understanding the effect of those factors on plant performance is required to predict the species responses to environmental change. We used reciprocal gardens planted with distinct regional *Andropogon gerardii* ecotypes adapted to dry, mesic, and wet environments to characterize their rhizosphere communities using 16S rRNA metabarcode sequencing. Even though the local microbial pool was the main driver of these rhizosphere communities, the significant plant ecotype effect highlighted active microbial recruitment in the rhizosphere driven by ecotype or plant genetic background. Our data also suggest that ecotypes were more successful in recruiting rhizosphere community members unique to their local homesites, supporting the “home field advantage” hypothesis. These unique homesite microbes may represent microbial specialists that are linked to plant stress responses. Further, our data support ecotypic variation in the recruitment of congeneric but distinct bacterial variants, highlighting the nuanced effects of plant ecotypes on the rhizosphere microbiome recruitment. Our results should facilitate expanded studies on understanding the complexity of plant host interactions with local soil microbes and identification of functional potential of recruited microbes. Our study has the potential to aid in predicting ecosystem responses to climate change and the impact of management on restoration practices.

**Importance:** In this study, we used reciprocal gardens located across a sharp precipitation gradient to characterize rhizosphere communities of distinct dry, mesic, and wet regional *Andropogon gerardii* ecotypes. We used16S rRNA amplicon sequencing and focused oligotyping analysis and showed that even though the location was the main driver of the microbial communities, ecotypes could potentially recruit distinct bacterial populations. We showed that different *A. gerardii* ecotypes were more successful in overall community recruitment and recruitment of microbes unique to the “home” environment, when growing at their “home site”. We found evidence for “home field advantage” interactions between the host and associated rhizobiomes, and the capability of ecotypes to recruit specialized microbes that were potentially linked to plant stress responses. Our study provides insights into the understanding of factors effecting the plant adaptation, improving management strategies, and predicting of the future landscape under the changing climate.

## Introduction

The rhizosphere is a dynamic region characterized by the complex interactions between the plant host and associated microbial communities [1,2]. It has been widely recognized that plants directly and indirectly benefit from associated microbial activities and resultant microbial compounds [3,4]. Complex microbe-microbe interactions around the rhizosphere result in facilitating nutrient transformations, uptake, and cycling as well as altering soil structure and soil water availability [5,6]. Similarly, plant host-microbe interactions can also greatly affect the overall plant health and productivity [7,8]. Root-associated microorganisms (hereafter referred to as rhizobiome) can affect plant resistance to biotic and abiotic stress, help with nutrient uptake and even alter plant morphology and phenology [9,10]. Given the crucial roles of the rhizobiome, understanding what shapes microbial community assembly, function, and mechanisms, as well as adaptive responses between plant host and associated microbes is critical in predicting the response of this system to changing environmental conditions [11,12].

Numerous studies have demonstrated the need to consider the interactive effects of the environment and plant host genetics in understanding the rhizobiome. The chemical and physical characteristics of the local soil can directly affect plant function, which in turn can have consequential influence on the rhizobiome composition [11,13,14]. Plant hosts actively modulate associated microbial communities by releasing various signaling molecules (phytohormones) and compounds into the soil [11,15]. Phytohormones are structurally diverse secondary metabolites released by the plant to activate the immune system [16,17] in response to microbial pathogens [18] and even insect herbivores [19]. In addition to direct immune responses, plants produce phytohormones and compounds in response to abiotic stress such as nutrient or water deficiency, or to promote symbiotic interactions with soil microbes [20]. Although there are numerous studies on plant host influence on rhizobiome composition, only little information is available about the interactive impact of the host-environment on the rhizobiome. The rhizobiome is a subset of microorganisms available in the surrounding soil microbial pool [15]. Thus, it is no surprise that the same plants perform differently in distinct locations. Some studies have described an increase in efficacy of the plant host-associated rhizobiome interaction due to the “*home field advantage*” [21,22]. However, questions remain if plants locally adapted to the prevailing environmental conditions can take advantage of the local soil microbes or if plants preferably favor microbiomes similar to their home environment.

Due to climate change, drought frequencies and severity are predicted to increase across the United States [23-25]. In the Great Plains, drought events limit productivity especially in tallgrass prairies [24]. Thus, understanding the effect of drought in shaping the rhizobiome and the mechanisms of mutual adaptation between plants and associated microbes is critical in the prediction of the ecosystem’s response to changing climate. To date, most studies [26-28] on the effect of drought on plant-microbe interactions have been under *in vitro* conditions using model organisms, lacking the complexity and dynamics of the natural system application [29,30].

To address the question if there is a co-adaptation of the plant host and its associated rhizobiome in a non-model plant system, our study took the opportunity to investigate the rhizobiome composition of three locally adapted big bluestem (*Andropogon gerardii* Vitman) ecotypes planted in reciprocal common gardens across the precipitation gradient in the Great Plains. *Andropogon gerardii* is a perennial C4 grass that is widely distributed across the Great Plains of North America [31,32], and covers up to 80% of the biomass in tallgrass prairie [32-35]. Within the Great Plains, *A. gerardii* has been growing for over 10,000 years along prominent sharp rainfall gradients that range from semiarid to heavy rainfall [36,37]. Time and environmental heterogeneity lent support for local adaptation of *A. gerardii*, giving rise to distinct ecotypes (dry, mesic, and wet) [36,38,39] [36,39,40]. Previous investigations have revealed that *A. gerardii* ecotypes vary in functional traits that influence microbially mediated processes [40,41]. Although there are studies investigating intraspecific [42] and interspecific (reviewed in [40]) plant responses to climate [43], the role of the rhizobiome in plant host adaptation in the natural system remains unclear.

Here, we (1) investigated the relative importance of the plant local environment and phenotypic variation of *A. gerardii* on establishing plant rhizobiome; (2) compared ecotypic responses in their ability to recruit rhizosphere microbes under reciprocally transplanted, local and non-local environments; and (3) examined rhizosphere community members with the potential to link the microorganism recruitment and the plant hosts’ ecotypic resilience functions. We hypothesized that *A. gerardii* ecotypes would perform better at the location closely matching their “home” environment, highlighting the effect of the plant genetic background or ecotype on rhizosphere community assembly. We also predicted that greater diversity or locally adapted unique microbes would be recruited in the homesites. Our work contributes to a clearer understanding of the factors that influence the plant host’s recruitment of rhizobiome and will help to provide insights into *A. gerardii* ecotypic responses to a future changing climate.

## Materials and Methods

### Study sites, samples collection and processing

We sampled three *A. gerardii* reciprocal gardens in the summer of 2019. The gardens had been established in 2009 and continually maintained at the three sites: Hays, KS (H) at Kansas State University Agriculture Experimental Station (38°85’N, 99°34’W), Manhattan, KS (MHK) at USDA Plant Materials Facility (39°19’N, 96°58’W), and Carbondale, Illinois (C) at Southern Illinois University Agriculture Research Station (37°73’N, 89°17’W), giving us an excellent opportunity to study the interactive effects of local environment and hosts on the rhizobiome (Supplementary Table S1). Experimental details of the reciprocal garden experiment have been published in Galliart et al. [44]. Briefly, in 2009, seeds were collected from four *A. gerardii* populations at each of three different locations (12 populations in total): mixed grass Central Kansas (CKS/Dry: CDB, REL, SAL, WEB), tallgrass Eastern Kansas (EKS/Mesic: CAR, KON, TAL, TOW) and Southern Illinois savanna (SIL/Wet: 12MI, DES, FUL, WAL) (Acronyms of populations are listed in Supplementary Table S1) prairies across the natural rainfall gradient with 580 mm/year, 871 mm/year and 1167 mm/year of mean annual precipitation, respectively. We defined the ecotypic variation of *A. gerardii* based on the locations they were collected (Dry: Hays; Mesic: Manhattan and Wet: Carbondale). Seeds representing the three ecotypes and twelve populations were germinated and grown in a greenhouse using potting mix (Metro-Mix 510). Established 3-to 4-months-old seedlings were then planted at each three reciprocal garden sites (size - 4 × 8 m), in which 12 plants (4 populations × 3 ecotypes) were planted in a complete randomized block design with 10 blocks (rows) for a total of 120 plants per site. Plants were planted 0.5 m apart along each row, and the soil around the plants was covered with the water-penetrable landscape cloth for weed control.

Some plants did not survive transplanting or through the ten years of growth in the common gardens. As a result, we were able to sample a total of 284 *A. gerardii* rhizospheres across the three locations (Supplementary Table S2) using a soil core (15 cm deep × 1.25 cm diameter). Samples were sealed in Ziplock bags, transported on ice, and stored at -20°C until DNA extraction.

We extracted total DNA from 0.150 g of rhizospheric roots and soil using an Omega E.Z.N.A. Soil DNA Kit (Omega Bio-Tek, Inc., Norcross, GA, USA) as per the manufacturer’s protocol with a slight modification. We removed soil from the plant roots by shaking; any soil that remained attached to the roots was considered rhizosphere soil [45,46]. We mechanically lysed the cells on a Qiagen TissueLyser II (Qiagen, Hilden, Germany) using glass beads for 2 mins at 20 rev/s prior to any downstream DNA extraction steps. The extracted DNA was eluted to 100 μ L final volume. The DNA yield and concentration was measured using a Nanodrop and a Qubit™ dsDNA BR Assay Kit. Extracted DNA was sequenced (2 × 250 cycles) using Illumina MiSeq with the 16S rRNA V4 region amplified using the primers 515F and 806R at the Kansas State University Integrated Genomics Facility.

### Sequence data processing and analyses

We used QIIME 2 v. 2021.4 [47] to process a total of 8,353,179 raw sequences, resulting in 5,628,314 bacterial sequences after quality control. We used QIIME 2 plugin cutadapt [48] to remove the primer sequences. Any sequences with ambiguous bases, with no primer, with greater than 0.1 error rate mismatch with primer or any mismatches to the sample-specific 12 bp molecular identifiers (MIDs) were discarded. Following initial quality control, we used DADA2 [49] with the same parameters across different runs. We used the pre-trained SILVA database (v. 138) in QIIME 2 for taxonomic assignment of the bacteria. We rarefied the data set to 10,000 reads per sample (resulting in 2,104,416 high quality sequences) to minimize biases resulting from differences in sequencing depth among samples before estimating diversity indices and downstream analyses [50].

We used Analysis of Variance (ANOVA) to test for main and interactive effects in observed richness (S_Obs_), community (Shannon’s _H’_), and phylogenetic (Faith’s PD) diversity of the rhizosphere associated bacterial communities among the locations (Hays (H), Carbondale (C), Manhattan (MHK)), ecotypes (dry, mesic, wet), and populations (12MI, CAR, CDB, DES, FUL, KON, REL, SAL, TAL, TOW, WAL, WEB) nested within ecotype. Following overall ANOVA, we used pairwise comparisons using Kruskal-Wallis test to identify factors that were driving the significant effects.

We estimated pairwise Bray-Curtis distances to compare the bacterial communities among the different factors and visualized these data using Non-Metric Multidimensional Scaling (NMDS) ordinations. We used a nonparametric permutational analog of traditional analysis of variance (PERMANOVA) to determine whether bacterial composition differed among locations, ecotypes, and populations. In this model, “population” was nested within “ecotype.” Following the PERMANOVA, we performed pairwise comparisons using pairwise.adonis function. We also used betadispr function to test whether the community dispersion differed between any significant groups. To identify bacterial populations that were disproportionately more abundant in one significant group than in another, we analyzed these data using indicator taxon analyses. The community composition data analyses were conducted in R ape [51], indicspecies [52], lme4 [53], phyloseq v 1.42.0 [54], stat v 4.1.1, vegan [55], R studio [56].

### Oligotyping analyses

We used minimum entropy decomposition (MED) [57] algorithm with default parameters to identify sequence variants in high-throughput 16S rRNA amplicon sequences. The MED partitions the sequences into discrete sequence groups by minimizing the total entropy in the dataset. We then concatenated the sequences assigned to a specific genus using silva 138 reference database and used the supervised oligotyping method available in the oligotyping pipeline version 2.1 [58]. In this supervised oligotyping method, we used Shannon entropy, with a threshold value of minimum 0.2 and minimum substantive abundance threshold of 10, to obtain genus-level oligotypes. We selected *Pseudomonas* and *Rhizobium* oligotypes for further analysis due to the overall high relative abundance across the bacterial ASVs and generated oligotypes (Oligos: *Pseudomonas*-Oligos (n=6); *Rhizobium*-Oligos (n=6))

## Results and discussion

We analyzed a total of 2,104,416 (19,131 ± 4,115 per sample) high quality sequences assigned to 1,157 ASVs. Of the recovered reads, an average of 90% was annotated to the genus level on Naive Bayes classifier. Any unknown or unclassified ASVs were removed from downstream analyses. The bacterial taxon assignments along with their counts in each sample are provided in Supplementary Table S3. Our data were dominated by the phylum Proteobacteria (45% of all sequences), followed by Actinobacteria (15%), Acidobacteria (12%) and Bacteroidata (6%). A small proportion of sequences remained unclassified (0.29%). ASV relative abundances were dominated by Proteobacteria (28% of all ASVs), Actinobacteria (14%), Firmicutes (12%) and Bacteroidetes (9%). Similar to sequences, the proportion of unassigned ASVs was small (0.3%).

### Location had a strong impact on the ecotypic rhizobiome richness and diversity

Location was the main driver of the bacterial communities, with a strong effect on bacterial richness and diversity (ANOVA, S_Obs_F_2,245_=7.13; P < 0.001; Shannon’s H’ index: F_2,245_= 26.56; P < 0.001; Faith’s PD index: F_2,245_ = 4.09; P = 0.018) (Figure 1A). Pairwise tests showed that community richness (S_Obs_) and Shannon’s diversity were lower in Manhattan than in the other two locations (Table 1). On the other hand, the phylogenetic diversity (Faith’s PD) in Carbondale was marginally higher than in Hays and Manhattan (Table 1). Phylogenetic diversity analyses suggest that Manhattan was favorable for generalist microbes - fast growing and abundant soil microbes that are good at colonizing the plant and often characterized by the lower diversity [2,4] due to the intermediate environmental conditions as compared to Hays and Carbondale. The higher bacterial phylogenetic diversity we observed in this study could be highly attributed to the amount of precipitation among the locations.

**Figure 1.**
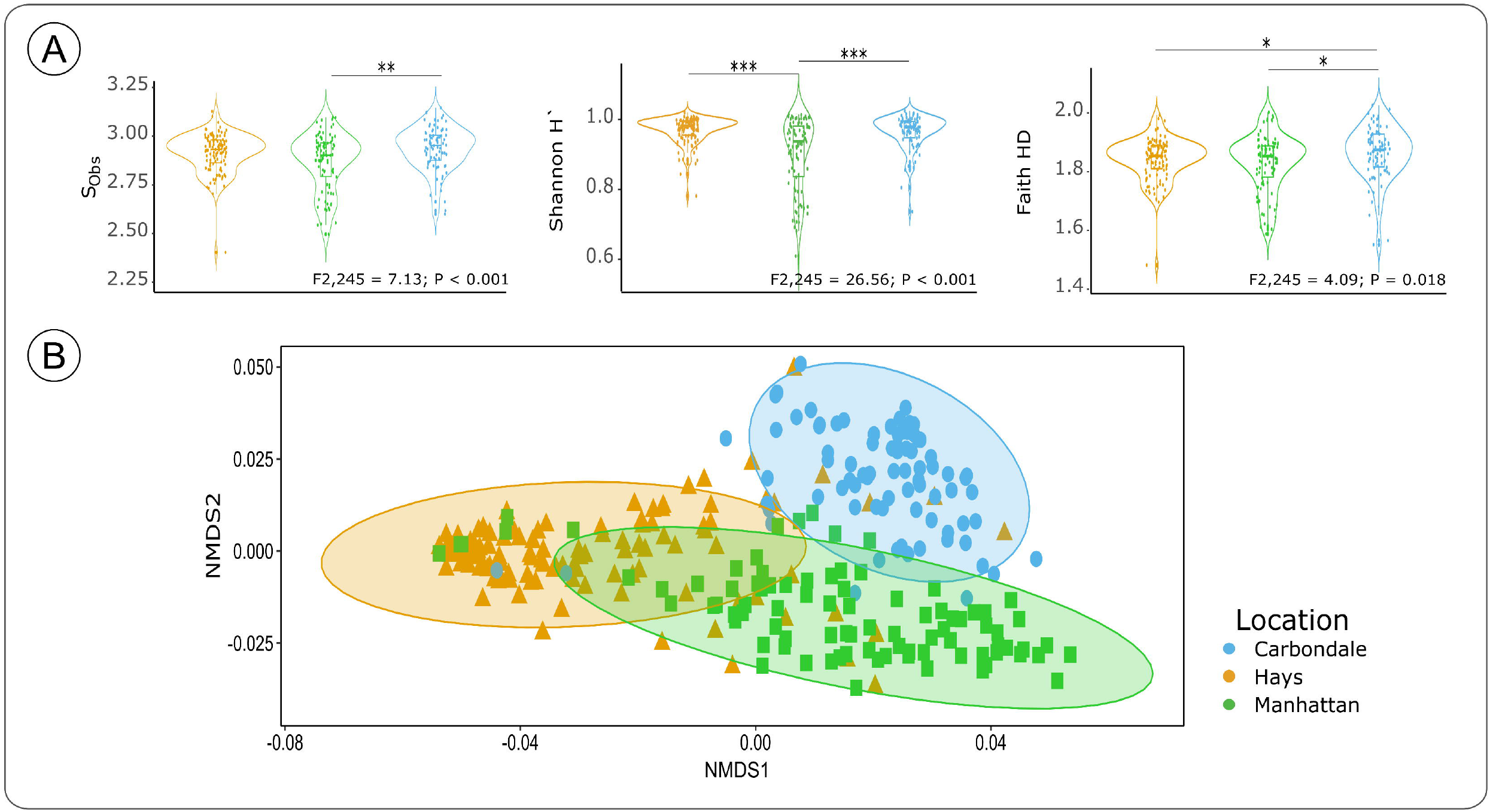
(A) Bacterial α-diversity indices among three sites located across precipitation gradient: Hays, Manhattan, and Carbondale. Species observed richness (S_Obs_), Shannon’s H index, and Faith-PD index. (B) Location impact on *Andropogon gerardii* rhizosphere bacterial composition. Non-metric multidimensional (NMDS) scaling plot of *A. gerardii* rhizosphere communities associated with three site locations across the precipitation gradient: Hays, Manhattan, and Carbondale. NMDS ordinations were obtained from Bray-Curtis similarity matrix. (p-values: * - 0.05, ** - 0.01, *** - <0.01).

**Table 1.** Statistical analyses (ANOVA and pairwise test) reveal locations (C-Carbondale, H-Hays, MHK-Manhattan) had a strong influence on the rhizobiome diversity and richness.

| Index | ANOVA |  | Pairwise test |  |  |
| --- | --- | --- | --- | --- | --- |
|  | chi-squared | P-value | C | H |  |
| SObs | 10.42 | 0.005 | 0.1005 | - | H |
|  |  |  | 0.0067 | 0.0653 | MHK |
| Shannon's H | 25.171 | 3.422e-o6 | 0.7 | - | H |
|  |  |  | 6.90E-05 | 1.40E-05 | MHK |
| Faith's PD | 6.7829 | 0.03366 | 0.044 | - | H |
|  |  |  | 0.044 | 0.585 | MHK |
|  | chi-squared | P-value | Dry | Mesic |  |
| SObs | 3.46 | 0.033 | 0.585 | - | Mesic |
|  |  |  | 0.078 | 0.026 | Wet |
| Shannon's H | 0.91 | 0.404 | 0.285 | - | Mesic |
|  |  |  | 0.285 | 0.061 | Wet |
| Faith's PD | 2.51 | 0.0108 | 0.78 | - | Mesic |
|  |  |  | 0.14 | 0.13 | Wet |

Similar to the alpha-diversity, the bacterial communities clustered by location indicate that locations had strong effects on bacterial composition (PERMANOVA, R^2^ = 0.42; F_2,245_ = 102.53; P = 0.001; Figure 1B). Our pairwise comparisons further corroborated that all locations had significantly different bacterial compositions (Pairwise Adonis, C vs H: R^2^ = 0.435; P = 0.001; P_adj_= 0.003; C vs MHK: R^2^ = 0.313; P = 0.001; P_adj_= 0.003; H vs MHK: R^2^ = 0.274; P = 0.001; P_adj_=0.003). We conducted a dispersion analysis, and observed that bacterial communities in Manhattan (F_2,279_= 3.899; P = 0.015) were more varied than Carbondale (P = 0.020). Similar to richness, the variation in bacterial community compositions among the locations were likely a result of precipitation-associated biotic and abiotic differences [59] as well as interspecific microbial competition [2,4]. For example, the mean precipitation in Carbondale is 1167 mm/yr which is drastically different from that in Hays (580 mm/yr). This precipitation gradient among the 3 locations would result in differences in pH which is known to be a key factor influencing bacterial diversity [60,61]. In this study, we showed that location had a strong effect on bacterial diversity and composition. We next address if there are any bacterial populations that were disproportionately higher in relative abundance among the locations.

Our indicator taxon analyses identified 194 taxa that differed among the locations. The majority of the taxa were associated with the wettest and driest locations (Carbondale (n=181); Hays (n=151)), whereas fewer were associated with the mesic intermediate location [Manhattan (n=38)) (Supplementary Table S4), suggesting that Carbondale and Hays harbored higher number of habitat specialist microbes due to the prevailing environmental conditions [62,63]. Of our three common gardens, Manhattan had the most intermediate conditions for growth of *A. gerardii*, where plants were less likely to experience abiotic stress including water availability stress in Hays and waterlogged conditions in Carbondale. We further observed that the majority of soil bacterial indicator ASVs were assigned to the following four phyla - Acidobacteria (Total n=31; C = 21, H = 5, MHK = 5); Actinobacteria (Total n=66; C = 15, H = 47, MHK = 4); Planctomycetes (Total n=20; C = 13, H = 7, MHK = 0); and Proteobacteria (Total n=109; C = 53, H = 41, MHK = 15); and Verrucomicrobia (Total n=23; C = 14, H = 9, MHK = 0) (Supplementary Table S4). It is unsurprising that these phyla dominate in our indicator taxon analysis since they are ubiquitous among rhizosphere associated bacterial communities [41,64-66]. However, these indicators were not evenly distributed across locations, suggesting that this pattern was due to the soil moisture affecting the bacterial distribution. Soil moisture can directly affect the soil microbial composition and functionality by differentiating drought tolerance among taxonomic and functional groups of microorganisms [67,68]. For example, limited soil moisture restricts the solute mobility and therefore decreases substrate supply to the soil microbes. Therefore, highly moisture-sensitive Gram-negative bacteria populations (Acidobacteria, Planctomycetes, Verrucomicrobia) were affected by the limited water, resulting in lower number indicators in Hays and disproportionately higher in Carbondale [69-71]. In line with this argument, we also observed that Gram-positive bacteria (Actinobacteria) were disproportionately more abundant in Hays than in Carbondale and Manhattan [72].

### Ecotypes shaped the rhizosphere inhibiting bacterial richness and diversity

While there was a strong location effect on all alpha diversity metrics, a marginal ecotype effect was observed only in (S_Obs_) richness (ANOVA, S_Obs:_F_2,245_= 3.46; P = 0.033). The pairwise comparisons among ecotypes indicated that the mesic ecotype rhizobiome had a higher observed richness (S_Obs_) than the wet ecotype (Mesic vs Wet: H = 6.86, P_adj_= 0.026). We observed no differences in bacterial diversity and phylogenetic diversity among host ecotypes (ANOVA, Shannon’s H’ index: F_2,245_ = 0.91; P = 0.404; Faith’s PD index: F_2,245_ = 2.51; P = 0.108) (Figure 2A). It was not surprising that the effect of ecotype was lower than the effect of location. It is well discussed across the literature that the plant ecotypic effects can often be suppressed by the local biotic and abiotic environmental factors. For example, the environmental gradient defines the local microbial source pool available to the plant and therefore directly affects the source rhizosphere communities [18,73-75]. Taking into consideration the overwhelming effect of the local environment (location), we aimed to test if our ecotypes differed in bacterial richness (S_Obs_) locally. To do this, we split our data set by locations, and performed separate ANOVAs for each location with ecotype, population, and their interaction as the explanatory factors. Following that, we observed that only in Hays (F_2,87_= 15.06; P < 0.001), but not in Manhattan (F_2,87_= 0.10; P = 0.905) or Carbondale (F_2,79_ = 0.41; P = 0.668) did the ecotypes differ. We further used pairwise tests and identified that the wet ecotype in Hays had significantly lower richness (S_Obs_) compared to dry (Wet vs Dry: P_adj_ < 0.001) and mesic (Wet vs Mesic: P_adj_ < 0.001) ecotypes. These results highlight the potential differences in ecotype recruitment and structuring of the rhizosphere communities even when planted in a common environment. The overall effect of ecotype on bacterial community composition was also weaker than the location effect in the overall model (PERMANOVA, R^2^ = 0.01; F_2,245_ = 2.91; P = 0.012). We used pairwise comparisons and showed that the difference among the ecotypic bacterial composition was driven by the wet (Wet vs Dry: R^2^ = 0.028; P_adj_ = 0.015) and mesic ecotype (Wet vs Mesic: R^2^ = 0.038; P_adj_ = 0.015; Figure 2B).

**Figure 2.**
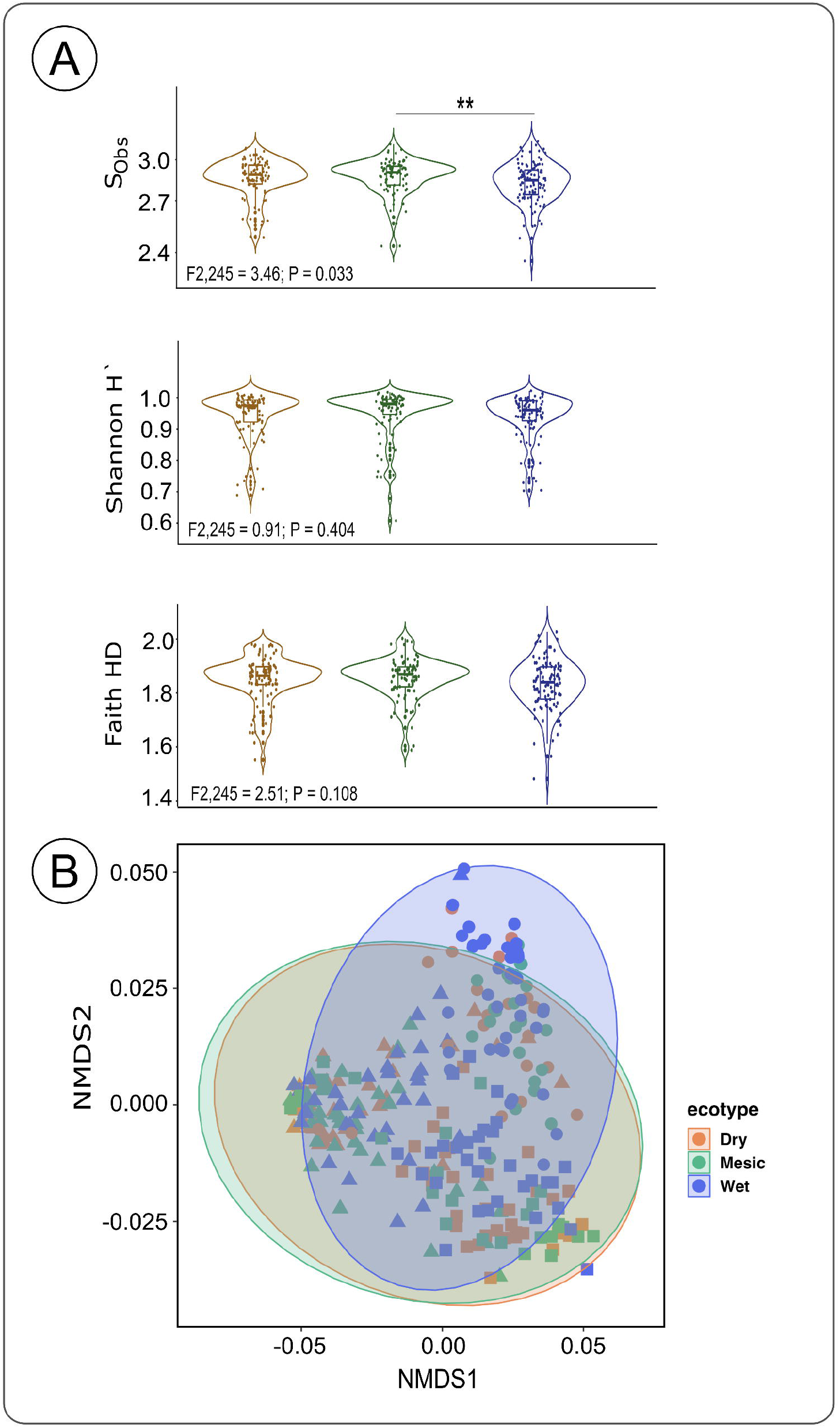
(A) Bacterial α-diversity indices among the dry, mesic, and wet ecotypes across all locations Species observed richness (S_Obs_), Shannon’s H index, Faith-PD index. (B) Ecotype impact on *Andropogon gerardii* rhizosphere bacterial composition. Non-metric multidimensional (NMDS) scaling plot of *A. gerardii* rhizosphere communities associated with three ecotypes across three site locations. NMDS ordinations were obtained from Bray-Curtis similarity matrix. (p-values: * - 0.05, ** - 0.01, *** - <0.01).

When we considered the impact of ecotypes on the unique microbial populations at each location, interestingly, we noticed that dry and wet ecotypes harbored more unique taxa when they were planted at their “*home field*”, except for mesic ecotype planted in Manhattan (C: Dry = 79, Mesic = 39, Wet = 127), (H: Dry = 121, Mesic = 71, Wet = 49) (MHK: Dry = 77, Mesic = 67, Wet = 72) (Figure 3C). Putting together our results on microbial community and indicator taxon analyses, we surmised that interaction between plant ecotypes and locations resulted in the wet and dry ecotypes harboring a higher number of habitat specialist microbes in Carbondale and Hays [62, 63]. While the dry ecotype was well-suited to its native environment in Hays, the wet ecotype’ rhizobiome differed suggesting that the wet ecotypic rhizobiome was not well-adapted to an environment (Hays) that is extremely different from its native location (Carbondale). Therefore, with a plant-host-rhizobiome mismatch in Hays, the wet ecotype would not be able to get a “*home field advantage*”, resulting in a higher number of generalists - abundant soil microbes that are good at colonizing plants [2,4,63,76,77]. Another limiting factor for wet ecotype to recruit and retain native specialist microbes in Hays is driven by lower drought stress tolerance of the plant host [36]. Wet ecotype is physiologically poorly adapted to the drier environment of Hays - which results in the lower photosynthetic rate [36]. We surmise that lesser volume of photosynthetic products exuded in the soil by the wet ecotype in Hays would limit the microbial taxa the wet ecotype could support in the drier environment. In addition, the limited photosynthates exuded by the wet ecotype would then be metabolized by the more abundant and faster colonizing generalists [63]. On the other hand, the dry ecotype in Hays produced lesser biomass but maintained higher photosynthetic rates, resulting in continuous and steady supply of photosynthate in limited quantities favoring slower growing microbial specialists [63]. Due to the general lower relative abundance of specialists in the samples, statistical analysis on ecotypic Shannon’s diversity and Faith PD as well as indicator taxon might not be sensitive enough to show the impact of ecotypes on its associated rhizobiome [78-80]. In order to test that idea, we calculated how many microbial taxa were uniquely observed in each location, to provide insights into the environmental and functional selection of potential specialists microbial populations. We showed that Carbondale (133) harbored the highest number of unique microbial taxa, followed by Hays (120) and Manhattan (79) (Figure 3A, Supplementary Table S5). We then looked at the unique taxa associated with ecotypes across all locations. Wet ecotype had the highest number of unique microbes (107), followed by Dry (92) and Mesic (66) (Figure 3B, Supplementary Table S5). Our results highlighted that ecotypes gained a “*home field advantage*” when planted in their native environment, and were able to match the plant host ecotypes to recruit and retain a higher number of unique microbial populations. On the other hand, the mesic ecotype was consistently associated with a lower number of unique microbial taxa. We hypothesize that the intermediate environment for *A. gerardii* in Manhattan resulted in harboring general microbial taxa that could be equally associated with all the ecotypes, however lacking the unique microbial drivers. Therefore, mesic ecotype was less adapted to recruit microbial specialists, and was associated with a lower number of unique taxa across all the locations.

**Figure 3.**
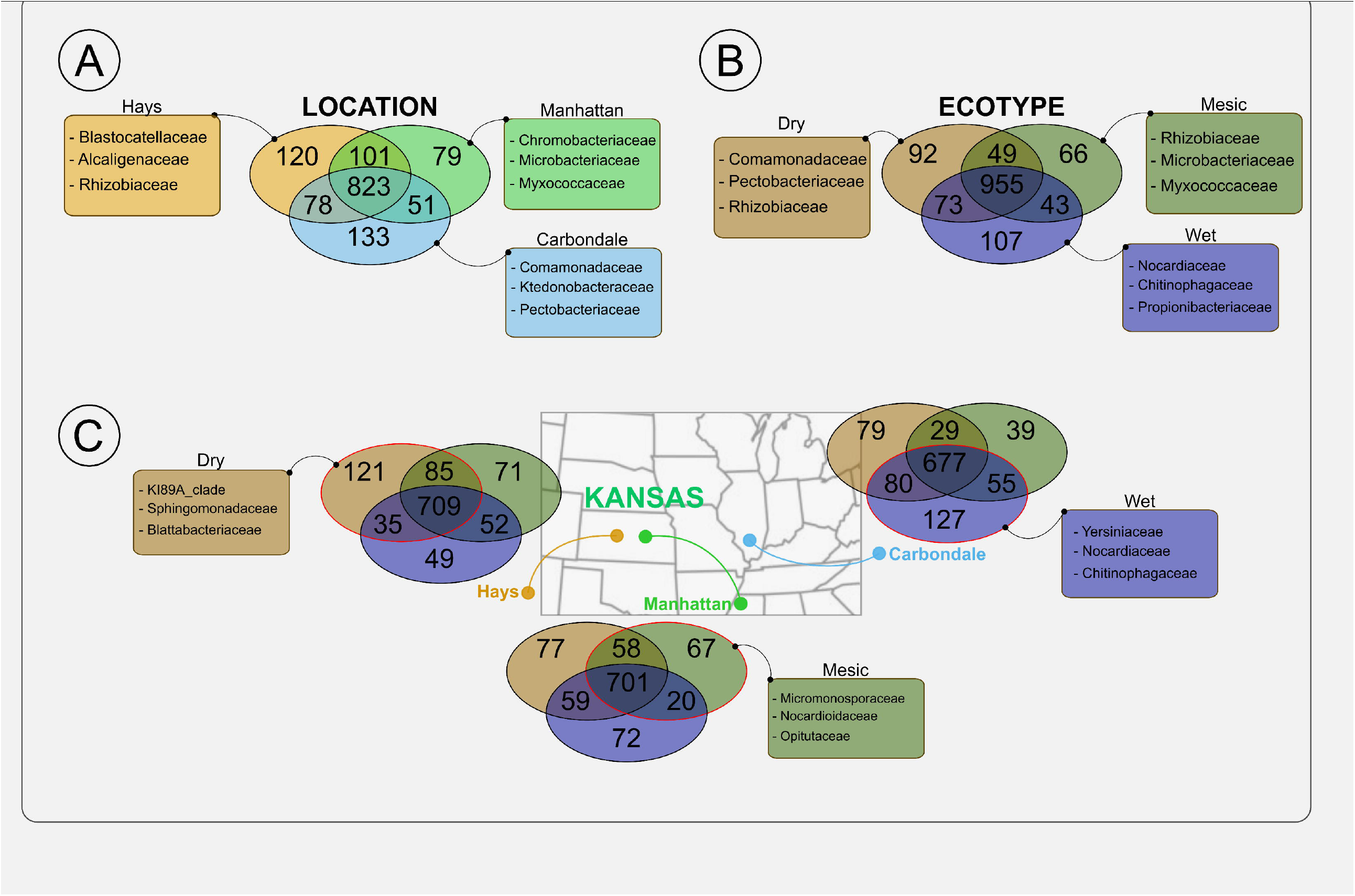
Venn diagrams representing the overlapping and unique ASVs (numbers in the circle) among (A) all locations, (B) all ecotypes, (C) ecotypes in Carbondale, ecotypes in Hays, ecotypes in Manhattan. The circle with a red edge represents the home ecotype for that location. Inserts represent the top three genera. Partial information on unique ASVs are shown here, full information is in Supplementary Table S5. Higher numbers of unique ASVs suggest the “*home field advantage*” of the *Andropogon gerardii* ecotypes.

### Matching of the plant host ecotype and rhizobiome was evident at the ecotype home location

We used indicator taxon analysis to give insights into the interactive relationship between the ecotypic plant host and its associated rhizobiome (Supplementary Table S6). We identified numerous indicator taxa that were significantly different among the ecotypes. We further showed that, considering individual locations, there was a strong effect of the ecotypes on the indicator taxa. Our study demonstrated that plant host ecotypes matched best with their rhizobiome, showing the highest number of indicator taxa, when the plant ecotypes were at their “*home field”*.

In the indicator taxon analysis across all the locations, we identified 26 indicator taxa that differed among the ecotypes. Wet ecotype was associated with the highest number of indicators (n=20), followed by mesic (n=6) and dry (n=0) ecotypes (Supplementary Table S6). The majority of soil bacterial indicator ASVs were assigned to Acidobacteria (Total n=10; Dry = 0, Mesic = 2, Wet = 8) and Proteobacteria (Total n=10; Dry = 0, Mesic = 1, Wet = 9). Similarly, taking into consideration the strong effects of location, we further focused our analysis by studying the ecotypic effects at different locations. We observed that while ecotypes differed at all locations (PERMANOVA, H: F_2,109_ = 2.55; P = 0.021, MHK: F_2,89_ = 2.87; P = 0.014, IL:F_2,81_ = 2.55; P = 0.015) (Figure 3C), in our pairwise comparisons of community composition in Carbondale and Manhattan, wet ecotype only differed from mesic (C: Wet vs Mesic: P_adj_ = 0.033, MHK: Wet vs Mesic: P_adj_ = 0.042), while other ecotype comparisons across all locations were not significant (P_adj_ > 0.05).

In Carbondale, we further observed that even though the ecotypic rhizobiome dispersion were significantly different (F_2,79_= 3.84; P = 0.026), dry ecotypic rhizobiome was more dispersed than those of mesic and wet ecotypes. In addition, we did not identify ecotype-associated indicator taxa in Carbondale among the ecotypes (n=0), highlighting that the differences in ecotypic recruitment might be on a smaller scale of unique and lower abundance taxa (Supplementary Table S7). In Manhattan, mesic ecotype was significantly more dispersed (F_2,87_ = 10.72; P = 0.01) when compared to dry and wet ecotypes. In Manhattan, we identified 11 indicator taxa - 4 indicators were associated with mesic (Actinobacteria=2; Proteobacteria=1; Verrucomicrobia=2) and 7 indicators with wet ecotypes (Actinobacteria=3; Proteobacteria=3; Cyanobacteria = 1). We observed that in Hays, wet ecotype differed in composition from dry and mesic ecotypes (Dry vs Wet: P=0.046, Mesic vs Wet: P=0.030), but no significant differences were observed after adjusting for multiple comparisons (P_adj_ =0.05). Interestingly, ecotypes did not differ in Hays (P_adj_ =0.05). Hays is at the very edge of continuous distribution of where *A. gerardii* are commonly found [36], suggesting that the drier environment might have a strong impact on diversity of bacterial populations that can proliferate under the arid conditions [81, 82]. Therefore, in Hays, we hypothesize that *A. gerardii* ecotypes 1) had a lower diversity of bacterial populations for recruitment; 2) experienced high abiotic stress from the drier conditions; and 3) had to compete for the recruitment of the smaller microbial specialist populations. Although our results suggest that all ecotypes had a more challenging survivorship in the harsher environment, will the dry and wet *A. gerardii* ecotypes be more resilient and successful in recruiting microbes due to their “*memory*” from surviving in a harsh “*home*” environment [21,83]? Will ecotypes get a “*home field advantage*” in the recruitment of microbes at their “*home*” locations?

To test that, we performed the indicator taxon analysis on ecotypes at each location (Supplementary Table S8). Dry ecotype recruited 64 indicator taxa ASVs in Hays, 43 in Carbondale and only 15 in Manhattan. Similarly, wet ecotype in Carbondale has the highest number of indicators (120), followed by Hays (76) and Manhattan (19). Mesic ecotype recruited 87 indicators in Carbondale, 51 in Hays and only 7 in Manhattan. Most of the indicator taxa from dry ecotype were Proteobacteria (41), Actinobacteria (32), followed by Acidobacteria (12). On the other hand, wet ecotype had disproportionately higher relative abundance in Proteobacteria (69), Actinobacteria (37), followed by Verrucomicrobia (19). Even though lower microbial recruitment in Manhattan could be generally due to lower number of indicators in Manhattan as we discussed earlier, it is clear that both wet and dry ecotypes recruited more indicator taxa when they were growing in their home environments, suggesting how “*home field*” confer a physiological and adaptive advantage to the plant host and associated rhizobiome.

Since we observed differences in indicator taxa and unique ASVs across locations and ecotypes, we further conducted oligotyping analysis to inspect the concealed diversity within ASVs at a higher resolution. (Supplementary Table S9). We noticed that the relative abundance of ASVs and oligos associated with *Pseudomonas* and *Rhizobium* displayed opposite abundance patterns across the locations (Figure 4). Pseudomonadales had lower relative abundance of ASVs across samples from Hays and Carbondale compared to samples from Manhattan (Figure 4A). On the other hand, the relative abundance of Rhizobiales was similar throughout Hays and Carbondale and had a high degree of differences between samples in Manhattan (Figure 4B). Interestingly, we noticed dissimilarities among locations and ecotypes across both *Pseudomonas-*Oligos and *Rhizobium-*Oligos (Figure 4). Even though the relative abundance distribution patterns of oligos was obviously driven by location, we observed differences in ecotypic recruitment across ecotypes. Further work is necessary to provide more insights into recruitment patterns of strain-level Pseudomonadales and Rhizobiales populations by the plant host to understand the impact of bacterial populations on hosts’ resilience to environmental stresses. However, our results suggested that precipitation-induced stress (drier in Hays and wetter in Carbondale) could result in a shift of plant host-rhizobiome interaction, changing the *Pseudomonas*-Oligos and *Rhizobium*-Oligos composition. For example, in Hays, *Pseudomonas*-Oligos-GG were more prominent in dry ecotype as compared to wet ecotype, while *Pseudomonas*-Oligos-AG were highly detected in mesic ecotype (Figure 4A). On the other hand, in Manhattan, *Pseudomonas*-Oligos-GA decreased in relative abundance in both dry and wet ecotypes in Manhattan, with an increase in relative abundance of *Pseudomonas*-Oligos-AA. Similarly, in Carbondale, there was a higher relative abundance of *Pseudomonas*-Oligos-AA wet ecotype as compared to dry ecotype (Figure 4A). We further noticed that in Hays, there was a higher relative abundance of *Rhizobium*-Oligos-ATATC in dry ecotype as compared to wet ecotype. However, the relative abundance of *Rhizobium*-Oligos-ATATC was higher in the wet ecotype as compared to dry ecotype in Manhattan. Interesting *Rhizobium*-Oligos-TGATG was substantially higher in relative abundance in wet and mesic ecotypes as compared to dry ecotype in Carbondale (Figure 4B). The differences in the *Pseudomonas*-Oligos and *Rhizobium*-Oligos across the locations and ecotypes potentially support the “*home field advantage*” and tendency of plants to recruit microbial strains that could contribute to the plant hosts’ ecotypic resilience [21,84]. The intraspecific genome content may be particularly important in understanding these host-microbe interactions.

**Figure 4.**
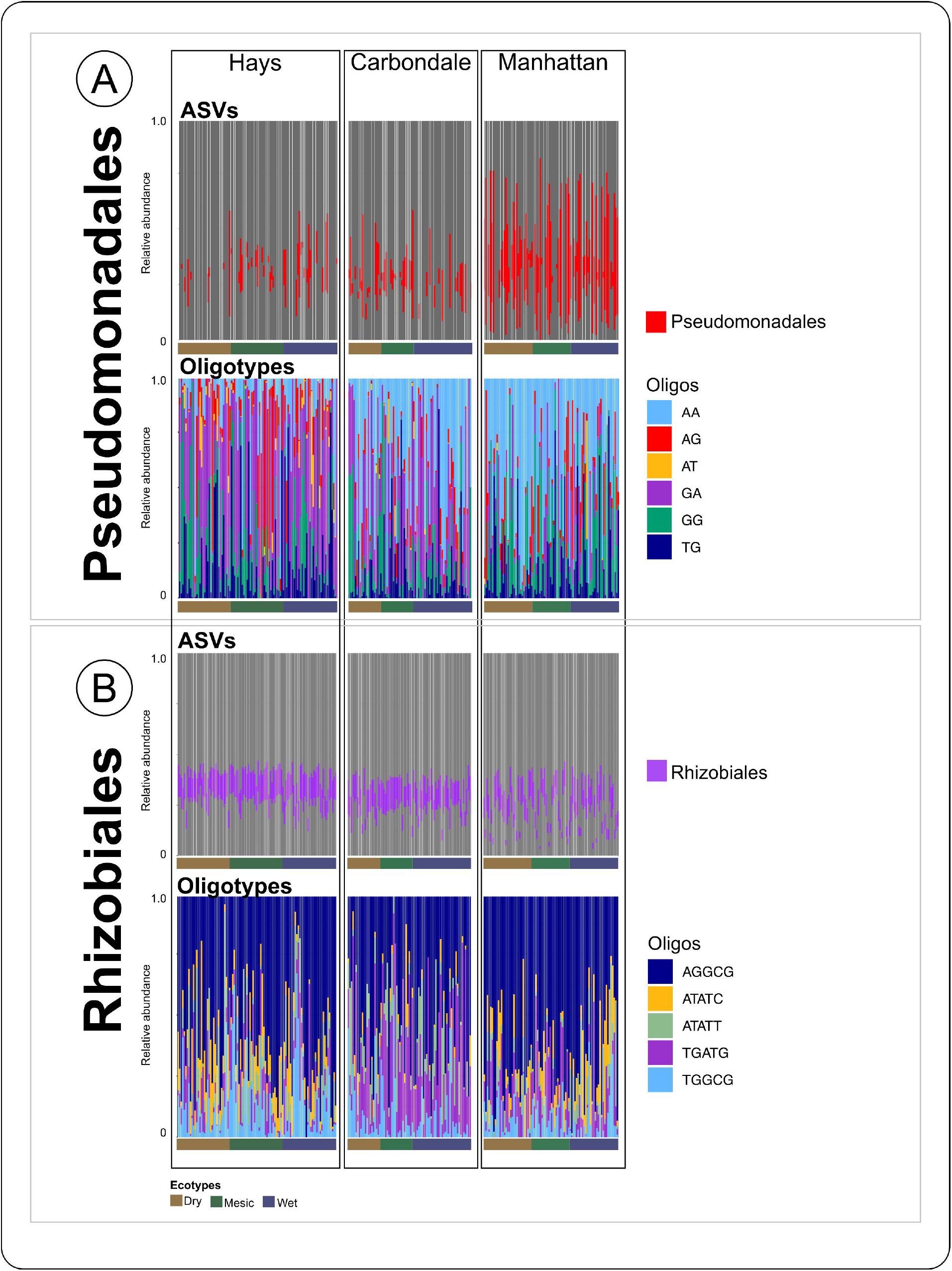
Relative abundance of each oligotype within the Pseudomonadales and Rhizobiales diversity for each sample among three sites located across precipitation gradient. The proportion of the relative abundance of the (A) Pseudomonadales (red, top) and *Pseudomonas*-Oligos (bottom), and (B) Rhizobiales (purple) and *Rhizobium*-Oligos (bottom) suggest the impact of locations and ecotypes on rhizobiome strain-level composition.

## Conclusion

Plant responses and adaptation to stress depend on a combination of environmental factors and plant genetics [1,2]. While the environment defines the soil-associated microbial pool, our results highlighted the plant-host’s ability to recruit soil microbial communities under different environmental conditions. We showed that plant hosts were more successful in the recruitment of both the general and unique microbial population when grown at their homesite [85]. Our observations also suggested the “*home field advantage*” and mutual association between plant and specific soil microbes [21,22,84]. From our study, we further propose that, at their home sites, ecotypes originating from these harsher environments, experiencing constant abiotic stresses, were better able to recruit specialist microbes with the potential stress relief functions [85]. We argue that in the study of plant host-rhizobiome, in line with the generalist and specialist hypothesis, plant host resiliency under abiotic stress are more dependent on specialized microbes [2,4,63,76,77] rather than the core microbiome [15,86-88], Our study suggests that in terms of the resistance to abiotic stress, the key factor is in less abundant and specialized microbes with specific functions rather than general common microbes which are abundant in soil. We further observed, in our study, the fragile relationships between plant hosts and associated specialized microbes. Although it might be challenging to tease apart the “*specialist vs generalists*” and “*home field advantage*” concepts, we used oligotyping analysis and showed the ecotypic variations in recruitment of bacterial stains from the same genera. These observations even further highlight the complexity of these symbiotic relationships. After growing in a common garden for over 10 years, we could still see the distinct microbial communities recruited by the ecotypes. However, the disconnect between the plant host ecotypes and rhizobiome in the harsher environment of Hays demonstrated that the plant host-rhizobiome interaction is much more fragile than we imagined. Taking all these factors in consideration, there is a crucial gap to identify specialist microbes, and understanding these relationships and communication signals with plant hosts, to predict the response of species to the changing climate change.

## Data availability

The raw sequence data are available through the Sequence Read Archive under BioProjectBioSamples PRJNA911775, while bioinformatic and statistical codes are available through figshare (10.6084/m9.figshare.21791306).

## Acknowledgements

This work was supported by the National Science Foundation EPSCoR Award No. OIA-1656006 and matching support from the State of Kansas Board of Regents. This study was supported by the United States Department of Agriculture, National Institute of Food and Agriculture (USDA NIFA), under the Award Number: 2020-67019-3180. We would like to thank Sara Baer for aiding in the designing of the reciprocal gardens. We would like to thank all of those who assisted in sampling and data acquisitions. We are grateful for the help of Tanner Richie, Brandi Feehan with the discussion on bioinformatics and data analysis. We are grateful for the technical assistance by Alina Akhunova, Sarah Bastian and Samantha Elledge at the Integrated Genomic Facility at Kansas State University (https://www.k-state.edu/igenomics/).

## Author Contributions

A.J., L.J. and S.T.M.L. designed the study. L.J. designed the reciprocal gardens experimental design, and A.J., L.J. and S.T.M.L. maintained sites. Sample collection was performed by S.S., E.H., M.G., A.J., S.T.M.L. and S.T.. S.S., A.K., K.W. extracted DNA with Nanodrop and Qubit quality analysis. A.K. and Q.R. performed 16S rRNA sequencing bioinformatic analysis. Q.R. and S.T.M.L generated oligotypes data. A.K. performed statistical analyses. A.K. and S.T.M.L wrote the manuscript, prepared figures, and supplementary files. A.J. and L.J. authors aided in manuscript and figure revisions. S.T.M.L., A.J. and L.J.acquired fundings for this study. All authors read, contributed to manuscript revision, and approved the submitted version.

## Conflict of Interest

The authors disclose no conflicts of interest.

## Supplementary Tables

Supplementary Table S1. Locations of seeds and populations collected from *Andropogon gerardii*.

Supplementary Table S2. Samples names and details for the soil cores collected for this study.

Supplementary Table S3. Raw sequence analysis by QIIME 2 Version 2019.7. The number of counts of bacteria and fungi initially obtained, and the counts that were considered after primer trimming and DADA2 quality control per sample. Bacterial identifications along with the number of counts per sample.

Supplementary Table S4. Indicator taxon analysis on different locations (irrespective of ecotypes) - Hays, Manhattan, and Carbondale.

Supplementary Table S5. Unique taxa for locations and ecotypes.

Supplementary Table S6. Indicator taxon analysis on ecotypes (irrespective of locations) - Dry, Mesic, and Wet.

Supplementary Table S7. Indicator taxon analysis on locations by individual ecotypes.

Supplementary Table S8. Indicator taxon analysis on ecotypes by individual locations.

Supplementary Table S9. Genus-level oligotyping analysis.

## Notes

https://figshare.com/search?q=10.6084%2Fm9.figshare.21791306

https://www.ncbi.nlm.nih.gov/bioproject?term=PRJNA911775

### Competing Interest Statement

The authors have declared no competing interest.

### Summary of Updates

Importance section added Figure 3 revised Figure 4 revised Results and discussion of oligotyping analysis is revises Supplementary Table S1 updated

